# Integrated, high-dimensional analysis of CD4 T cell epitope specificities and phenotypes reveals unexpected diversity in the response to *Mycobacterium tuberculosis*

**DOI:** 10.1101/2024.11.05.622086

**Authors:** Heather L Mead, Jung Hwa Kirschman, Caroline E Harms, Erin J Kelley, Jorge Soria-Bustos, Georgia A Nelson, Paul Ogongo, Gregory Ouma, Samuel G Ouma, Joel D Ernst, John A Altin

**Affiliations:** The Translational Genomics Research Institute (TGen), Flagstaff, AZ, USA; Gilead Sciences, Foster City, CA, USA; The Translational Genomics Research Institute (TGen), Phoenix, AZ, USA; Division of Experimental Medicine, University of California, San Francisco, CA, USA; Center for Global Health Research, Kenya Medical Research Institute, Kisumu, Kenya

## Abstract

Immunity to *Mycobacterium tuberculosis* (*Mtb*), like many pathogens, is encoded jointly by the antigen specificities and functions of responding CD4 T cells. However, these features span a large two-dimensional possibility space – defined on one axis by the *Mtb* proteome, and on the other by the T cell transcriptome – that exceeds the dimensionality of existing technologies. Here we present an approach (“CRESTA”) that combines highly-multiplexed DNA-barcoded epitope probes, single cell sequencing, and clonal analysis of T Cell Receptors (TCRs) to robustly detect rare antigen-specific CD4 T cells across hundreds of epitopes simultaneously and reveal their transcriptome-wide phenotypes. By comprehensively assaying known epitopes in *Mtb*-infected participants, we reveal polyclonal and multi-epitope responses across a spectrum of differentiation states, uncover previously-unobserved phenotypic diversity within and between epitopes, and increase the total number of known *Mtb* epitope-mapped TCRα:βs by ∼8-fold. We expect CRESTA to enable high-dimensional analyses of CD4 T cell responses in various settings, including infection, cancer, autoimmunity and allergy.

## Introduction

Helper T cell immunity resides in populations of CD4+ cells that have clonally expanded in response to particular MHC class II-bound peptide antigens, and differentiated to acquire specialized effector functions^1–4^. Although these features together theoretically encode an individual’s state of immunity, they are difficult to measure in an integrated and comprehensive way because existing technologies do not scale well to the large diversity of possible CD4+ T cell antigen specificities and functions. Traditional assays for the identification of antigen-specific T cells include those that measure peptide-stimulated cytokine production (e.g., immunospot assays and flow cytometric detection of intracellular cytokines^5,6^) or upregulated markers^7^, as well as assays that use fluorescently- or isotopically-labeled peptide:MHC probes to directly detect antigen-binding T cells^8,9^. Although widely useful, these approaches have a limited ability to multiplex across antigens and phenotypic markers, meaning that available sample volumes and cell numbers quickly become limiting, typically precluding comprehensive analyses^10^.

More recently, DNA-barcoded peptide:MHC probes have been used to bypass the spectral limits of cytometric approaches (which generally peak at 10–50-plex), and have enabled CD8 T cell responses to be resolved across a multiplexity of up to ∼1000 distinct peptide:MHCs^11^. While it represents a major advance, this system has two key limitations: (i) by itself, it does not enable the simultaneous capture of T cell phenotypic information and (ii) it has not yet been adapted to the analysis of CD4 T cells. The latter likely reflects a combination of factors, including: (i) the lower frequencies of epitope-specific CD4 T cells, (ii) the generally lower affinity of CD4 T cell receptors (TCRs) for their MHC class II-restricted antigens, and (iii) greater challenges in identifying MHC class II:peptide binding pairs and constructing the corresponding probes. Indeed, despite considerable efforts to enable the multiplexed detection of CD4 T cell antigen-specificity ^9,12–18^, the greatest reported scale at which epitopes have been resolved simultaneously in a primary sample using any approach is 6-plex^19^. A powerful assay developed recently uses an elegant cell interaction reporter system to enable the genome-scale discovery of CD4 T cell antigens across 100,000s of candidate epitopes^20^, however it requires substantial genetic engineering of T cells and therefore does not enable the analysis of primary and/or polyclonal samples (currently it has been limited to the mapping of TCRs). Overcoming the barriers to highly-multiplexed analysis of CD4 T cell epitopes in primary samples would represent an important advance with widespread applications across the many settings in which helper T cell immunity is implicated.

The opportunities that would be enabled by high-dimensional assays of CD4 T cell specificity and phenotype are exemplified by *Mycobacterium tuberculosis* (*Mtb*), a pathogen responsible for 1.3 million global deaths in 2022, and for which an effective vaccine would have a major impact^21^. No vaccine for *Mtb* has been approved since BCG (∼100 years ago), although there are several experimental formulations under development^22–24^. A major challenge in the field has been selecting which *Mtb* antigens (from among >4000 proteins) to include in a vaccine, and what T cell functions it should elicit. Previous attempts to define protective immunity to *Mtb* have shown mixed success and have focused on diversity in one dimension at a time – eg using proteins / lysates to identify diverse CD4 T cell states (e.g. cytokine polyfunctionality^25,26^), or IFNγ production in response to diverse sets of peptides predicted to bind Human Leukocyte Antigen (HLA) proteins^27,28^.

More recently, sequence clustering approaches have been used to analyze the TCRs of T cells enriched for *Mtb* specificity and reveal public TCR groups^29,30^, including groups that are positively and negatively associated with disease progression^31^. Importantly, these findings indicate that particular antigen-specificities may be important in protection from *Mtb*. However, the analysis of TCRs alone does not enable the efficient identification of the cognate *Mtb* epitopes (which have been mapped for only a minority of TCR groups), nor does it capture the corresponding T cell phenotypes. These two missing features – antigen-specificities and phenotypes – are critical attributes if a correlate of protection is to be maximally actionable in guiding next-generation vaccine design.

Here, we present an assay that can measure reactivity across 100s of HLA class II:peptide antigen specificities simultaneously within a single PBMC sample, and associate each epitope-specific response with a transcriptome-wide T cell phenotype. We use this assay to generate a comprehensive portrait of the CD4 T cell response to *Mtb* in infected participants – molecularly resolved across 100s of epitopes and 1,000s of transcripts – and reveal previously undescribed polyclonality, phenotypic diversity and TCR sequence features within this response.

## Results

### Simultaneous, highly-multiplexed measurement of CD4 T cell epitope-specificity and transcriptional state enabled by clonal analysis

To enable the integrated, high-dimensional analysis of CD4 T cell epitope-specificities and transcriptional states, we adapted an approach developed previously for the highly-multiplexed analysis of CD8 T cell epitope-specificities^11^. That assay involved generating DNA-barcoded probes of MHC class I:peptide complexes multimerized onto dextran backbones, incubating T cells with pools of probes, using fluorescence to sort probe-binding cells, and analyzing binding by deep sequencing of the DNA barcodes. From this starting point, we introduced three main modifications, each of which was critical to enable our analysis of CD4 T cell responses. First, in place of MHC class I, we used MHC class II:peptide complexes, prepared by exchanging peptides of interest into HLA reagents bearing peptides tethered by a protease-cleavable linker ^32^. Second, we used single cell sequencing in place of bulk sequencing, to enable T cell transcriptional states to be detected and linked to epitope specificity. Third, instead of cell sorting, we introduced an antigen-specific clonal expansion step to overcome the low frequencies of circulating epitope-specific CD4 T cells, and then leveraged this expansion to boost analytical power by developing a “pseudobulk” approach in which we aggregated single cell data at the clonal level. Together, we refer to this workflow as the **C**lonally-**R**esolved **E**pitope-**S**pecificity and **T**ranscriptome **A**ssay (CRESTA) (**Figure 1**).

**Figure 1:**
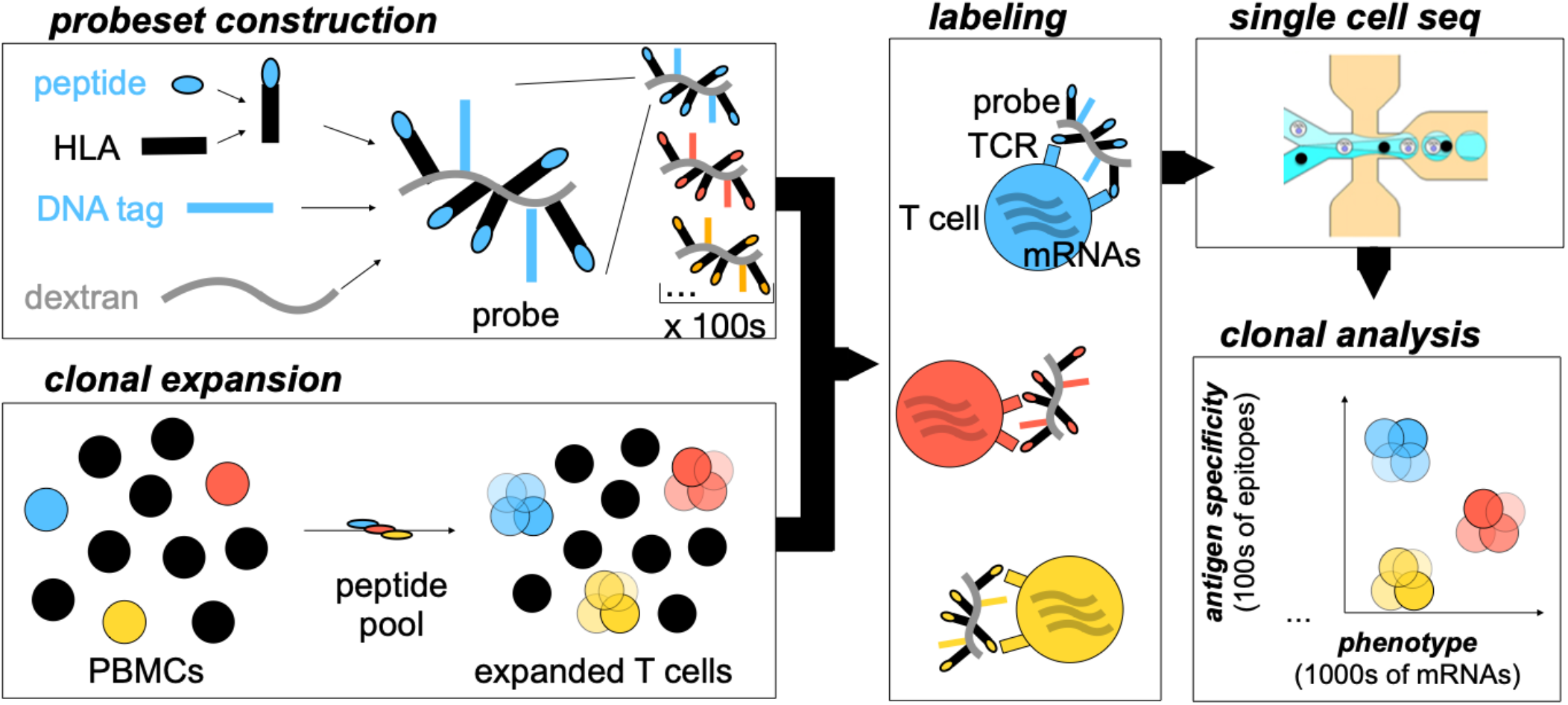
Workflow of the Clonally-Resolved Epitope Specificity and Transcriptome Assay (CRESTA). To enable the integrated, high-dimensional analysis of CD4 T cell epitope-specificities and transcriptional states, we begin with the high-throughput assembly of hundreds of peptide:MHC probes, each multimerized and DNA-barcoded using a streptavidin-bearing dextran backbone construct (*upper left*). Pools of these probes are used to stain T cells clonally expanded from PBMC samples using peptides (*lower left, center*), and then single cell sequencing is used to read out epitope-level specificities and transcriptome-wide expression profiles (*upper right*). Finally, to interpret the resulting data, we apply a “pseudobulk” analysis that aggregates cells into clones according to shared TCRα:β sequences (*lower right*), enabling inferences about epitope-specificity and transcriptional state that are robust to the noise inherent in single cell data.

We applied CRESTA to study the T cell response in HLA-typed (**Supplementary Table 1**), Quantiferon-positive, HIV-negative participants from a cohort in Western Kenya, sampled following recent (≤ 3 months) household exposure to active pulmonary tuberculosis^33^. To represent the known antigen space, we used the Immune Epitope DataBase (IEDB) to comprehensively identify a total of 206 *Mtb* peptide:HLA pairs known to generate human T cell responses in the context of 4 class II restriction elements that were prevalent in this cohort (HLA-DQB1^*^06:02, HLA-DRB1^*^15:03, HLA-DRB1^*^11:01, or HLA-DRB5^*^01:01). We also included 32 peptide binders for a fifth prevalent HLA – HLA-DRB1^*^01:02 – identified in an *in vitro* peptide:HLA binding assay. Finally, we selected a total of 85 additional peptide:HLA pairs as controls, which included CLIP-tethered (uncleaved) versions of each HLA, Mtb epitopes restricted by additional alleles (not expressed by the participants of interest), and IEDB epitopes from Influenza A virus (see **Supplementary Table 2** for a complete list of peptide:HLA pairs). We expanded antigen-specific T cells from 5 – 10 × 10^6^ PBMCs for 7-9 days using a master pool of all peptides. We then analyzed two aliquots of each sample: the first (“antigen-specificity aliquot”) was stained with pooled DNA-barcoded peptide:HLA class II multimers corresponding to the participants’ HLA types, the second (“gene expression aliquot”) was not stained but instead briefly restimulated with PMA/ionomycin to enhance antigen-responsive gene expression. This dual aliquot strategy was motivated by initial experiments (not shown) indicating that PMA/ionomycin greatly enhanced our ability to resolve T cell states, but led to diminished multimer binding, likely due to downregulation of the TCR^34^. Both aliquots of each sample were analyzed by single cell sequencing using the 10X Genomics Chromium platform to recover multimer barcode counts, transcriptome-wide mRNA abundance data and paired, full-length TCRα:β sequences on 1,000s-10,000s of single cells. We then informatically matched TCR sequences between the two aliquots to identify cells from common clones and thereby unite antigen-specificity and gene expression data for each.

In a representative participant (ID:40059), we recovered data on a total of 28,129 cells –17,312 and 10,817 for the antigen-specificity and gene expression aliquots, respectively – the former aliquot having been stained with a pool of 48 multimers (containing DRB1^*^15:03, DRB1^*^11:01 or DQB1^*^6:02, all expressed by the participant). We reasoned that TCRα:β sequences could be used to assign single cells into clonal families – each representing the progeny of a single *Mtb*-specific precursor – that each share a common epitope-specificity and differentiation state. To test this hypothesis, for each aliquot we organized individual cells into clones based on shared TCRα:β sequences, filtered on clones that contained ≥3 individual cells in each aliquot, and used a Kruskal-Wallis test to quantify the extent to which signal from peptide:HLA multimers and transcripts was driven by clonal identity (**Figure 2**). This analysis included a total of 170 clones, each of which contained 6 – 616 individual cells across the two aliquots (median=45) (**Figure 2a**).

**Figure 2:**
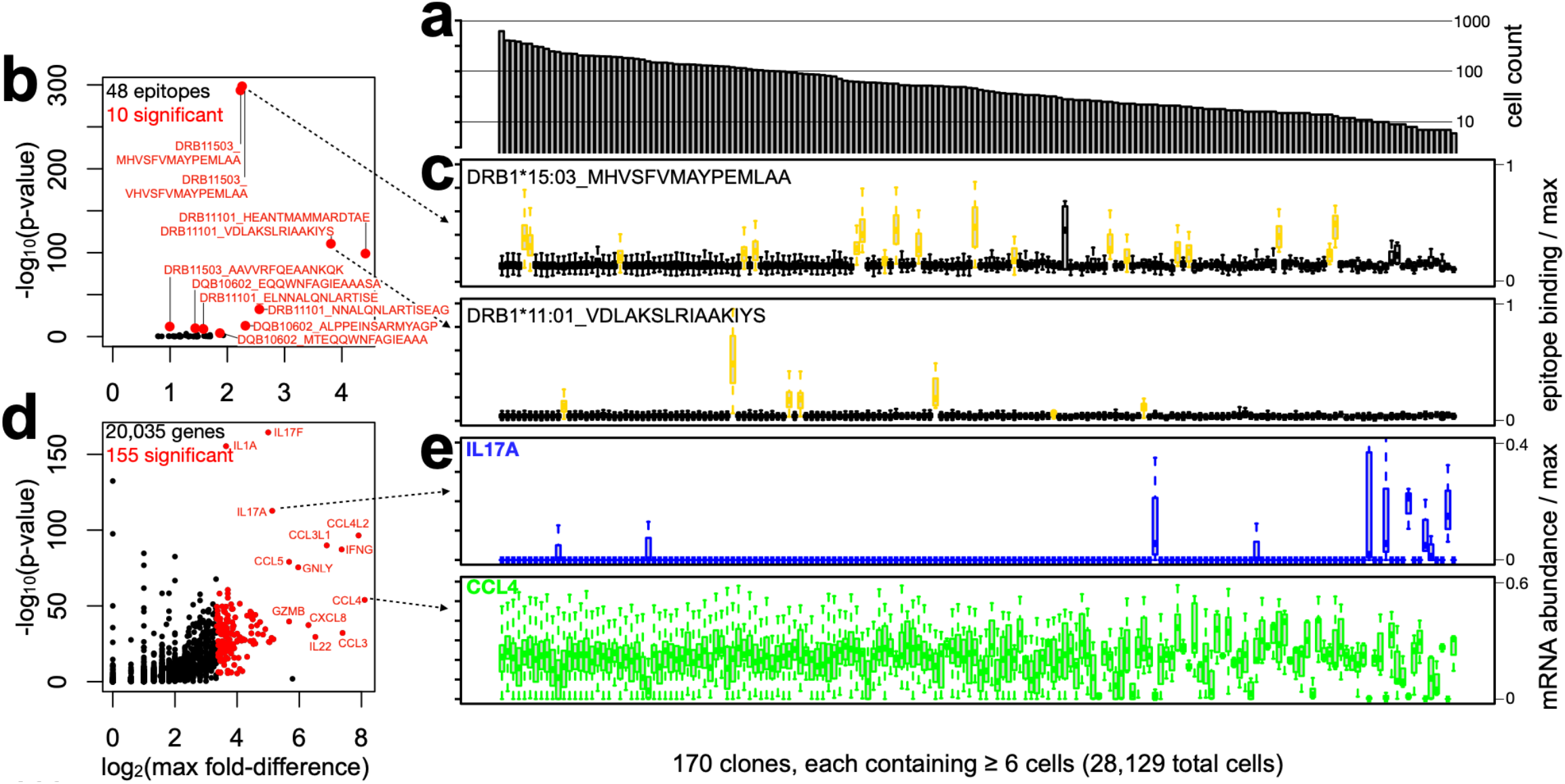
Clonal analysis enables robust, high-dimensional analysis of CD4 T cell *Mtb* epitope-specificity and transcriptome-wide gene expression, simultaneously. Peptide-expanded PBMCs from a LTBI participant (ID:40059) were stained with a pool of 48 DNA-barcoded peptide:MHC multimer probes corresponding to all *Mtb* T cell epitopes in IEDB known to be restricted by either HLA-DRB1^*^15:03, HLA-DRB1^*^11:01 or HLA-DQB1^*^6:02, and analyzed by single cell sequencing (“antigen-specificity aliquot”). A second aliquot that was unstained but stimulated with PMA/ionomycin was also assayed (“gene expression aliquot”). A total of 28,129 cells across the 2 aliquots were collapsed into clones based on identical TCRα:β sequences, 170 of which contained ≥3 cells in each aliquot (≥6 total). **(a)** Shown are the distribution of cell numbers in each of the 170 clones. **(b)** To identify which of the 48 epitopes were recognized, we applied a Kruskal-Wallis test to data from the “antigen-specificity aliquot” to determine whether, for each multimer probe, staining across cells partitioned in a clonally-restricted way (p-value, y-axis), and to quantify the magnitude of such partitioning (fold-difference, x-axis). **(c)** For significant epitopes, we next identified their particular binding clones using a Wilcoxon test to compare the distribution of probe binding to each clone against the distribution for all other clones. Shown, by way of example, are 2 of the significant epitopes identified in (b), with their respective significant clones indicated in yellow. **(d)** To identify genes whose expression varies by clone (“clonal differentiation genes / CDGs”), we applied the same clonally-resolved analysis described in (b), but this time to gene expression values measured in the “gene expression aliquot”. **(e)** Examples of the clonally-resolved expression patterns of 2 CDGs with anti-correlated abundance, corresponding to the Th17 (IL17A) and Th1 (CCL4) states, respectively. Vertical alignment of the 220 clones is consistent between panels (a), (c) and (e).

Consistent with the hypothesis, we observed strong partitioning of multimer binding by clone. In particular, we identified 10 of the 48 multimers to be binding in this participant (p-values ranging from ∼1e-5 to 1e-298), of which 3, 4 and 3 were restricted by DRB1^*^15:03, DRB1^*^11:01 and DQB1^*^6:02, respectively (**Figure 2b**). To identify the individual binding clones for each of these epitopes, we applied one-tailed Wilcoxon tests post-hoc to each binding multimer, in which we compared the binding of each individual clone to the overall distribution (**Figure 2c**). This analysis identified a total of 54 binding clones (32% of all clones), ranging from 1-19 per multimer, revealing extensive polyclonality in the response to individual *Mtb* epitopes. Binding clones were non-overlapping between multimers, with the exception of 3 pairs of multimers that featured overlapping peptides restricted by the same HLA and shared overlapping patterns of clonal recognition (discussed further below), supporting the fidelity of the process. Moreover, the TCR sequences of epitope-binding clones showed significant homology within epitopes and recapitulated known public motifs, as discussed further below.

Similarly, we observed strong partitioning of transcript abundance across clonotypes: applied to the same set of 170 clones, the Kruskal-Wallis test detected 155 genes at a Bonferroni-corrected threshold of p<0.05 and fold-difference threshold of 10 (**Figure 2d**). These genes, which we hereafter refer to as Clonal Differentiation Genes (CDGs), were strongly enriched for genes implicated in helper T cell function, and prominently featured cytokines and chemokines known to be produced by particular subsets, including IFNG, IL17A/F, IL1A, CCL3, CCL4, CCL5, GNLY, CXCL8, CCL3L1, CCL4L2, CCL3 and IL22. We also observed patterns that were consistent with the maintenance of distinct differentiation states across clones, for example the anti-correlated expression of the Th1 chemokine CCL4 and the Th17 cytokine IL17A (**Figure 2e**). Together, these results establish CRESTA as a method for robustly resolving antigen-specific CD4 T cells across many antigen-specificities simultaneously from a single PBMC sample, and associating each with a transcriptome-wide gene expression state.

### Comprehensive, hypothesis-free phenotyping of the *Mtb*-specific response using CRESTA

To perform a deep, multi-participant analysis of the epitope-specific CD4 T cell response to *Mtb* infection, we performed CRESTA on a total of 5 LTBI individuals (ID:40059, analyzed above, and 4 additional participants: IDs: 30128, 30133, 30129, 30168) from the Kenyan cohort. We stained cells from each participant with multiplexed probesets matching their HLA types (32-173 multimers per participant, across 5 distinct HLA class II alleles), and recovered single cell sequencing data on a total of 106,673 cells (4,400-14,496 and 7,866-13,475 per participant for the antigen-specificity and gene expression aliquots, respectively). In total, this yielded 1,496 evaluable clones (169–454 per participant).

Consistent with our observations above, epitope staining (analyzed as described in **Figure 2b**) partitioned strongly by clonal identity and revealed polyclonal responses across all participants. The highest dimensional staining was 173-plex in participant ID:30168, in which we observed robust binding across 15 distinct epitopes (**Figure 3a**), with a total of 126 epitope-binding clones. Analysis of these profiles revealed 7 epitope clusters (each containing 1-5 epitopes) within which there was extensive sharing of binding clones (**Figure 3b**). All 7 clusters were internally HLA matched and comprised sets of overlapping peptide sequences, indicative of a common core peptide epitope in each (but with possible, clone-specific additional contributions from flanking/polymorphic residues). Importantly, no clones were positive for epitopes from more than 1 cluster, indicating high assay specificity.

**Figure 3.**
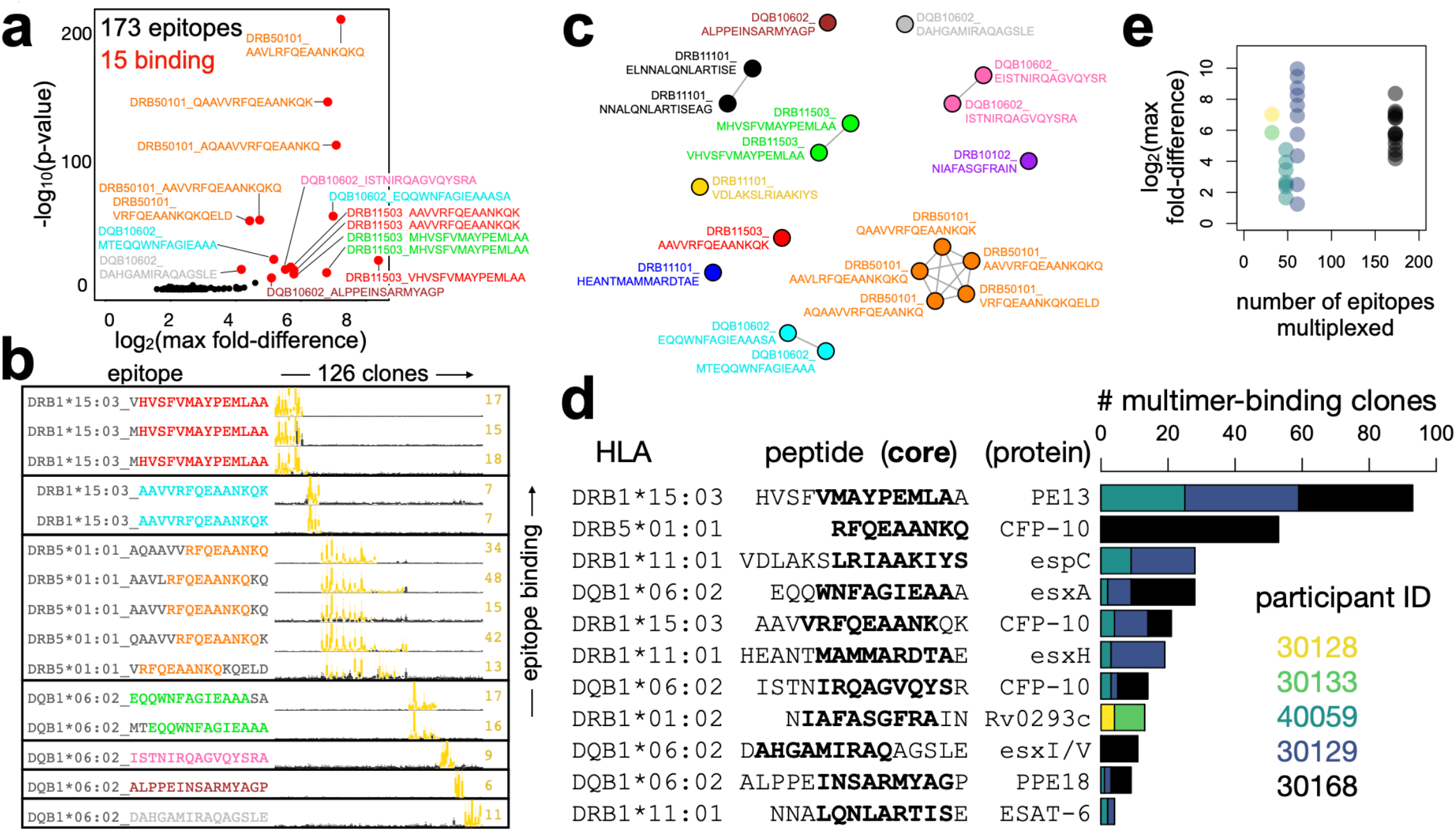
CRESTA epitope detection is specific up to at least 173-plex and reveals highly-polyclonal, multi-epitope responses to *Mtb*. **(a)** LTBI+ participant ID:30168 was analyzed as described in Figure 2a-c, this time using a pool of 173 DNA-barcoded multimer probes restricted by HLA-DRB1^*^15:03, HLA-DRB5^*^01:01 or HLA-DQB1^*^6:02. Shown is a volcano plot revealing the significant binding of 15 probes. **(b)** For the analysis in (a), boxplots show all 126 (of 454 total) clones for which we detected ≥1 significant probe binding event (highlighted in yellow) across the 15 probes. Based on these binding profiles, the 15 probes cluster into 7 groups (demarcated by horizontal boxes), whose members were uniformly HLA matched and contained overlapping peptide sequences (in 2 cases these groups contained identical replicates). Notably, none of the 126 clones showed significant staining across >1 of these groups (100% specificity on this measure). **(c)** The assay and analysis described in (a, b) was applied to a total of 5 participants (IDs: 30128, 30133, 40059, 30129, 30168), which revealed a total of 19 unique probes that had significant binding to ≥1 clone in ≥1 participant. Each of these probes is depicted as a node in a force-directed graph that connects probes whose binding clones overlap. This analysis yielded a total of 11 clusters (distinguished by color) which again correspond precisely to sets of probes with shared HLA and overlapping peptide sequences. **(d)** Shown are the number of significant binding clones for each of the 5 participants and 11 epitope clusters described in (c). Relative to (c), peptide sequences are trimmed to show only the longest subsequence common to all members of each cluster, and bold type indicates the HLA-binding core 9mer predicted by netMHCIIpan. **(e)** For each binding probe across the 5 participants (colored as in (d), and plotted by participant on the x-axis), shown is the intensity of probe staining (y-axis), which shows no decline with increasing plexity.

Across all 5 participants, we detected a total of 19 epitopes that showed significant binding to ≥1 clone in ≥1 participant (**Figure 3c**). Based on analysis of shared clones, these epitopes partitioned into 11 clusters, each of which exclusively contained members sharing a common HLA and overlapping peptide sequences, and we again found no examples of clones binding across clusters. For simplicity, in all downstream analyses, we refer to these clusters as “epitopes”, even though some comprise signal from several multimer probes that overlap a common core epitope. In total, we detected 293 epitope-specific clones in the antigen specificity aliquot, ranging from 4– 139 per participant and 4–93 per epitope. (**Figure 3d**). This number already represents nearly an order of magnitude increase over the total number of TCRα:β sequence pairs mapped to individual HLA II-restricted Mycobacterial epitopes in all prior studies to date (36 TCRs identified in 5 studies from 2009-2023: IEDB, queried 6/29/2024). Finally, across the range of 32-173-plex staining, we saw no evidence that the intensity of multimer probe binding was inversely correlated with the number of probes pooled for any given participant (**Figure 3e**), indicating that we have yet to approach the limit of assay multiplexing.

Next, we used the clonally resolved expression analysis described above in **Figure 2d**, to identify CDGs from the “gene expression aliquot” from each participant, and again observed these to be strongly enriched for known CD4 T cell differentiation genes. For each CDG, we collapsed individual cell expression values into per-clone medians, and then used UMAP to generate a 2-dimensional representation of the CDG-wide expression profile of each clone. This allowed us to cluster clones according to gene expression in a hypothesis-free fashion (**Figure 4a** shows participant ID:30133 as an example), and revealed distinct clusters whose expression profiles corresponded to those of naive, Th1, Th2 and Th17 cells. Beyond canonical gene expression patterns (i.e. IL2RA–, IFNG+, IL4+, IL17+, in naive, Th1, Th2, Th17 cells, respectively), hypothesis-free correlation analysis across all CDGs revealed that these clusters were each characterized by broad state-specific expression patterns, which included previously-described genes (e.g. TNF, CCL3, CCL4, HOPX in Th1 clones; IL3, IL5, IL13, GATA3, IL17RB in Th2 clones; IL17F, IL22 RORC, CCL20, NR4A2 in Th17 clones) as well as genes not previously associated with these T cell subsets (e.g. PTGS2 in Th2 clones; TGIF1 in Th17 clones) (**Figure 4b**). We also observed clones with a Treg-like expression pattern, characterized by high expression of FOXP3 and CTLA4, although these did not form a distinct cluster.

**Figure 4:**
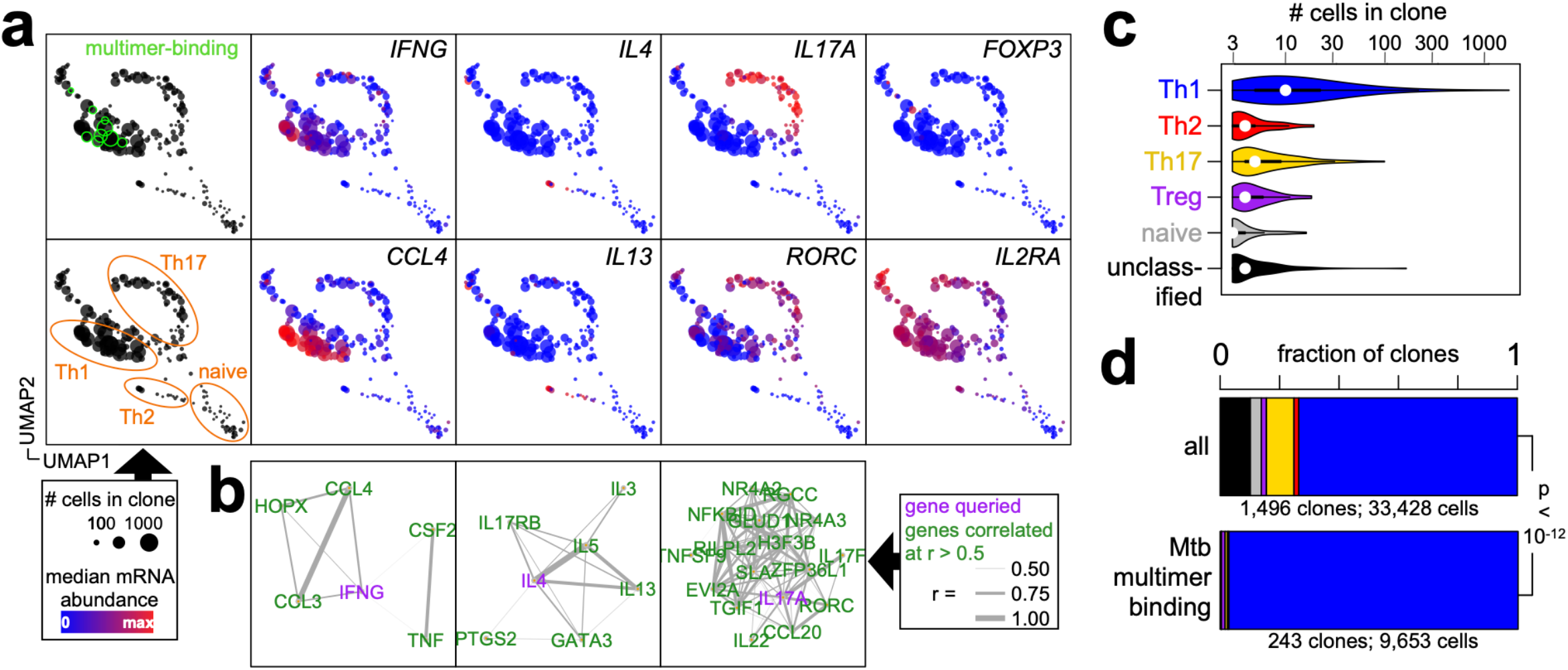
CRESTA reveals a spectrum of CD4 T cell differentiation states and localizes the epitope-specific *Mtb* response predominantly, but not exclusively, within the Th1 compartment. **(a)** Shown, for representative LTBI participant (ID:30133; 217 clones), is a clustering analysis in which each clone is represented as a circle (sized according to its number of constituent cells) and represented in 2 dimensions using a UMAP projection of its median expression of each Clonal Differentiation Gene (n=763 CDGs). Green circles (*upper left*) identify multimer-binding clones, orange ovals (*lower left*) demarcate inferred T cell subsets, and blue-red coloring represents the median clonal mRNA abundance for the genes indicated in the upper-right corner of each plot. **(b)** For the Th subset-specific cytokine genes IFNγ, IL4 and IL17A (represented in purple), Pearson analysis across all clones and CDGs was used to identify additional genes with correlated expression (represented in green). **(c)** Clones across 5 participants (n=1,496) were assigned to T-helper (Th) states Th1, Th2, Th17, Treg or naive states (or unassigned); the distribution of cell numbers per clone across the 6 Th states is shown. **(d)** The 1,496 clones described in (c) were further classified according to whether or not they bound an *Mtb* peptide:HLA multimer. Shown are the distributions across Th states of all clones (*upper*) or of multimer-binding clones (*lower*), which were compared by Fisher’s exact test.

Among a total of 1,496 clones analyzed for gene expression across the 5 participants, 1,344 (90%) could be assigned a differentiation state; composed of Th1 (74%), Th2 (2%), Th17 (9%), Treg (2%) or naive (4%) subsets. The average clone sizes across these diverse states differed significantly, with Th1 clones being the largest (median = 10 cells), followed by Th17, Th2, Treg and then naive clones (median = 3 cells) (**Figure 4c**). Of the 1,496 clones, we were able to assign *Mtb* antigen specificity to 243 (16%), across the 11 epitopes described above in **Figure 3** (50 of the 293 clones described in **Figure 3d** were not evaluable because although they had ≥3 cells in their respective antigen-specificity aliquot, they had <3 cells in the gene expression aliquot). Comparing these epitope-mapped clones to the total population, we observed a marked further skewing towards the Th1 state (97% of multimer-binding clones), at the expense of all other subsets (**Figure 4d**). As expected, no naive clones were found to be multimer binding. For this dataset, we therefore conclude that CD4 T cells specific for *Mtb*-specific epitopes are strongly biased towards the Th1 subset.

### Analysis of phenotypic heterogeneity within and between *Mtb* epitope-specific T cell responses

Despite the predominance of Th1 clones, we also detected rare *Mtb*-specific clones with Th17 (n=2) and Treg (n=2) phenotypes. A closer examination of these clones confirmed the expression of a range of transcripts corresponding to their respective states, and revealed that these patterns were broadly consistently across the individual constituent cells of each clone (**Figure 5a**). Notably, each of these clones existed alongside others specific for the same *Mtb* peptide:HLA epitope in the same participant, including the DRB1^*^15:03_HVSFVMAYPEMLAA (PE13) and DRB1^*^01:02_NIAFASGFRAIN (Rv0293c) epitopes. Together, this represents the first evidence of which we are aware that T cells recognizing the same *Mtb* epitope in the same individual can adopt highly-divergent differentiation states.

**Figure 5:**
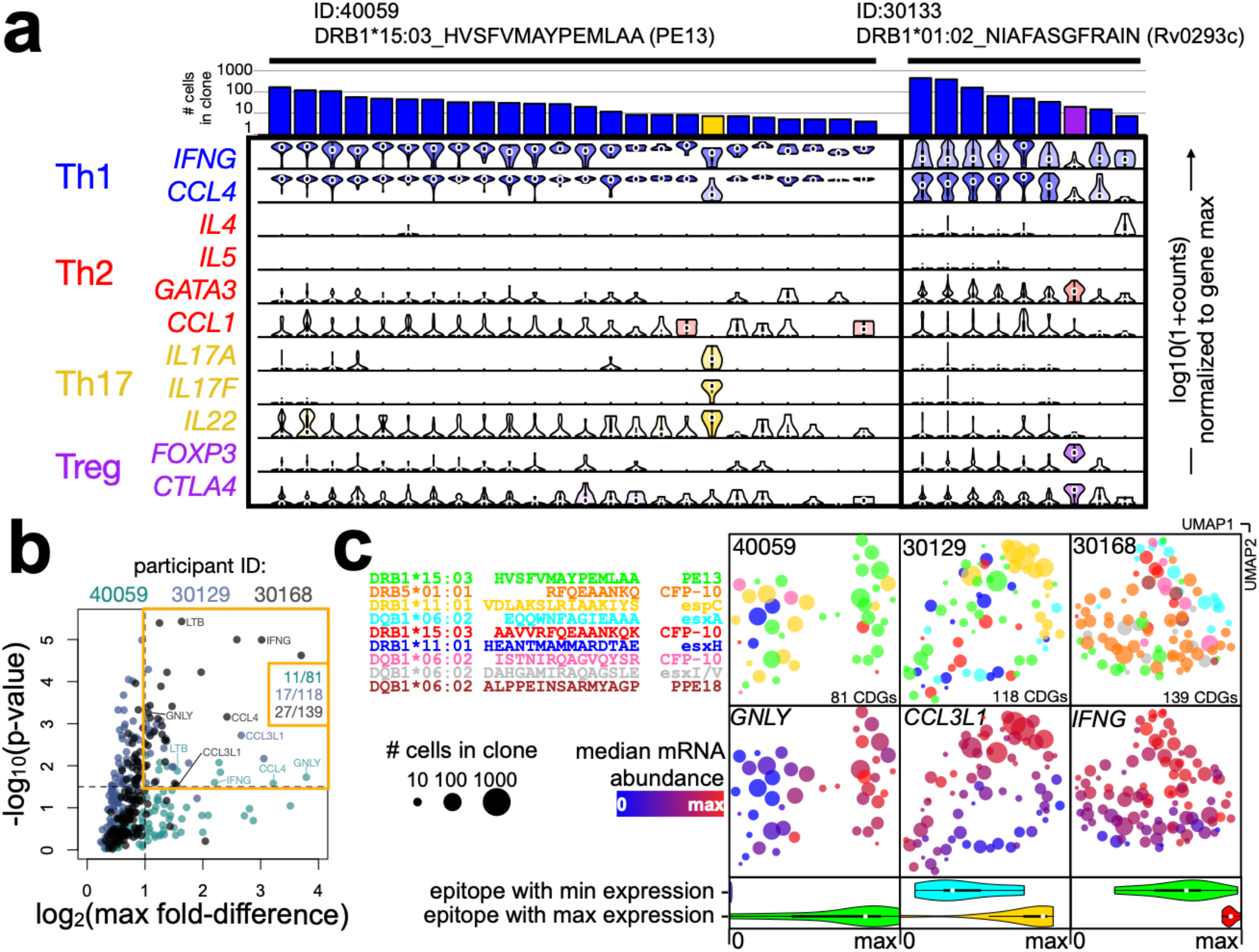
CRESTA reveals phenotypic heterogeneity within and between the CD4 T cell responses to individual *Mtb* epitopes. **(a)** Expression profiles of epitope-binding T cells for two selected individuals/epitopes, showing genes representative of the Th1 (IFNG, CCL4), Th2 (IL4, IL5, GATA3, CCL1), Th17 (IL17A, IL17F, IL22) and Treg (FOXP3, CTLA4) subsets. Each column shows data from an individual epitope-specific clone (n=33 clones), which comprise 4-347 individual cells, quantified in the barplot. Violin plots show the distribution of expression values for cells in each clone and are shaded according to median expression values. **(b)** For all cases in which we observed ≥2 distinct reactive epitopes each recognized by ≥3 clones within a participant (total = 230 clones across 9 epitopes in 3 participants), we identified CDGs within each participant and then applied Kruskal-Wallis tests to determine whether the expression of each CDG across clones was correlated with epitope specificity. Highlighted are 5 unique genes that were significant in ≥2 participants (using thresholds shown in the yellow box). **(c)** Gene expression states for the clones described in (b) were rendered in 2 dimensions using UMAP across all CDGs, and colored according to epitope specificity (dot plots, *upper row*). For each participant (column), expression of a selected gene observed to have strong epitope-dependent expression is shown across all clones (dot plots, *center row*), and compared between epitopes with the lowest v highest expression (violin plots, *lower row*). Violin plots are colored by epitope.

We next tested the hypothesis that, even within a Th1 dominated response, T cells recognizing different epitopes can have divergent gene expression programs. To address this in a systematic way, we considered all cases in which we observed ≥2 distinct reactive epitopes within a participant, each of which was recognized by ≥3 clones. These criteria focused the analysis to a total of 230 clones across 9 epitopes in 3 participants. We began by identifying CDGs as above, yielding a total of 81, 118 and 139 genes in participants ID:40059, ID:30129 and ID:30168, respectively. Focusing on these subsets of genes, we then used Kruskal-Wallis tests to compare the median gene expression of each clone across epitopes, measuring whether expression was correlated with epitope specificity. We detected strong correlations between gene expression and epitope specificity within all 3 participants, with p-values up to 1e-5 (**Figure 5b**). At thresholds of p<1e-1.5 and max fold difference >2, we identified 11, 17 and 27 epitope-linked genes in the 3 respective participants. These sets overlapped significantly: 5 genes were common to ≥2 participants – IFNG, CCL3L1, GNLY, CCL4, LTB – all of which have known roles in Th1 effector function.

To visualize how clones specific for different epitopes cluster in gene expression space, we used UMAP to render the CDG-wide expression profiles of each clone in 2 dimensions (**Figure 5c, upper row**). Rather than a random distribution of clones specific for different epitopes (indicated in different colors) across this space, in each participant we observed clustering of clones according to their epitope specificity, consistent with our Kruskal-Wallis analysis. Prominent among these were clusters of: (i) DRB1^*^15:03_HVSFVMAYPEMLAA-specific clones in participant ID:40059 enriched for GNLY expression (green dots in **Figure 5c**, upper left), (ii) DRB1^*^11:01_VDLAKSLRIAAKIYS-specific clones in participant ID:30129 enriched for CCL3L1 expression (yellow dots in **Figure 5c**, upper middle), and (iii) DQB1^*^06:02_EQQWNFAGIEAAA-, DRB1^*^15:03_AAVVRFQEAANKQK- and DRB1^*^15:03_HVSFVMAYPEMLAA-specific clones with high, high and low IFNG expression, respectively (cyan, red and green dots in **Figure 5c**, upper right). Together these results reveal that different epitopes can program diverse Th1 gene expression states within the same *Mtb* response, and that these are characterized by the differential expression of important effector genes.

### Analysis of TCR sequence clustering within the anti-*Mtb* response across epitopes and participants

An alternative approach, to the one described here, for the multiplexed detection of epitope-specific T cell responses has been the identification of clusters of homologous TCR sequences, followed in some cases by screening assays to map their specificities^30^. Applied to *Mtb*, this approach has been used to identify reactivities associated with protection from disease progression^31^. However, it remains unclear how TCRs identified by such homology-based approaches map onto the overall epitope-specific response.

The identification here of an unprecedented number of TCRα:β pairs mapped across a panel of *Mtb* peptide:HLA epitopes presented an opportunity to assess the degree of TCR sequence homology within epitope-specific T cell responses to *Mtb* in a minimally-biased way. To enable robust statistical power, we focused on the 8 HLA:peptide epitopes for which we detected ≥10 multimer-binding clones across participants – with the requirement that each clone have a single α and single β chain (which excluded 19% of clones, for which we sequenced 1 or >2 chains), to eliminate ambiguity about the chains involved in epitope binding – yielding a total of 206 TCRα:β pairs. We then used TCRdist^35^ to perform comprehensive pairwise sequence similarity measurements within each epitope. To rigorously quantify significance, we constructed a null distribution of >1e12 pairwise TCRdist measurements on a large set of TCRα:βs randomly-sampled from an unenriched repertoire. We used this distribution to transform TCRdist measures into p-values and applied Benjamini–Hochberg adjustment for the number of pairwise measurements performed for each epitope.

We detected significant TCR homologies for all 8 of the *Mtb* epitopes analyzed – evident both as the formation of clusters at an adjusted threshold of p<0.1 (**Figure 6a**), as well as overall deviations from the null p-value distribution (**Figure 6b**). However, the extent of this clustering differed markedly by epitope, encompassing up to 10/14 (71%) of TCRs specific for the esxH epitope DRB1^*^11:01_HEANTMAMMARDTAE, and as few as 2/13 (15%) for the CFP-10 epitope DQB1^*^06:02_ISTNIRQAGVQYSR. The PE13 epitope DRB1^*^15:03_HVSFVMAYPEMLAA, for which we detected the largest number of clustered clones overall, was notable for a large and unusually tight cluster of highly-homologous TCRs that was dominated by clones from two of the three participants that reacted to that epitope. For these TCRs, we observed a precise match to V/J segment usage (α: TRAV25/27, TRAJ52/40, β: TRBV9) and CDR3 motifs (α:CAG^***^S/TY, β:CASSVAL^*^G) described previously^30^, further supporting the fidelity of our multiplexed epitope-specific assay (**Figure 6c**). Overall, across the 8 epitopes, 99/206 (48%) epitope-specific TCRs were part of detectable homology clusters. Importantly, however, this degree of clustering was highly-dependent on filtering for single epitope binding to focus the number of TCR comparisons. When testing an oligoclonal scenario in which epitope-specific TCRs were diluted to 5% with random TCRs – designed to simulate more traditional enrichment methods where TCRs specific for >20 epitopes may be analyzed together – detectable clusters remained in only 5 of the 8 epitopes, and comprised just 24% of all TCRs.

**Figure 6:**
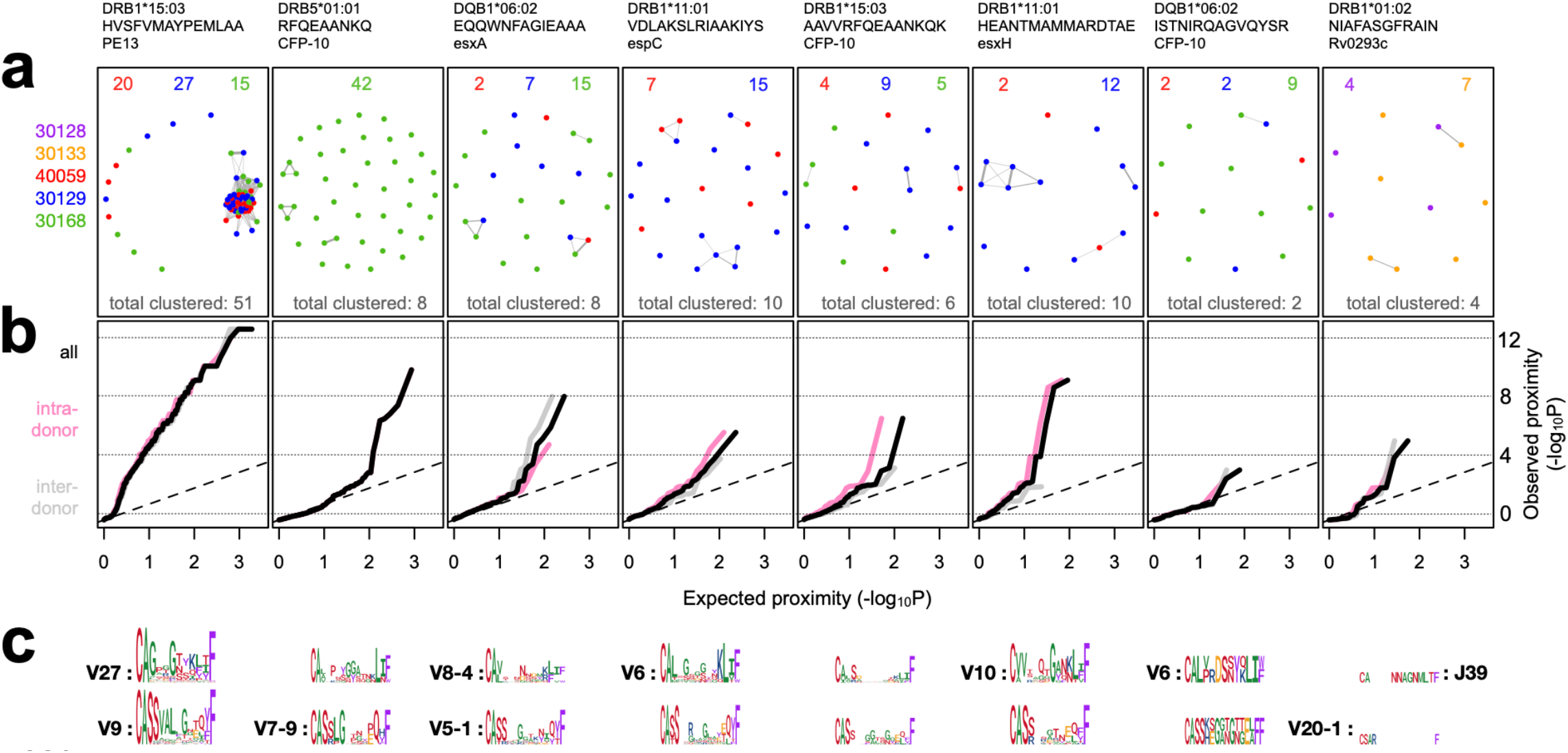
Analysis of 206 *Mtb* epitope-mapped TCRα:β pairs reveals that sequence publicity is widespread, but varies in magnitude across epitopes and participants. **(a)** For 8 *Mtb* epitopes (columns) for which CRESTA identified ≥10 epitope-specific clones, TCRα:β sequences were compared within and between participants using comprehensive pairwise TCRdist measurements. Each TCR is shown as a single dot (colored to distinguish participants) in a force-directed graph in which proximity indicates the degree of sequence similarity. Line segments connect TCR pairs whose TCRdist scores are significant, based on a large set of distances randomly-sampled from an unenriched repertoire, with Benjamini–Hochberg correction for the total number of comparisons made for each epitope (FDR<0.1). The number of epitope-specific TCRs for each participant, and the total number of TCRs within significant clusters, are shown at the top and bottom of each box, respectively. **(b)** Shown, for the analysis described in (a), are the full distributions of TCRdist p-values, but this time unadjusted: x-axis = expected, y-axis = observed), for all comparisons (black), and comparisons within (pink) and between (gray) participants. **(c)** Logos showing sequence features of the significantly-clustered TCRs for each epitope shown in (a). V and J segments are shown for cases where >50% of clustered TCRs contain the same segment. CDR3 letters are sized according to conservation/entropy, and colored by amino acid properties (polar=red, non-polar=green, negatively charged=gold, positively charged=blue, aromatic=purple).

Many of the observed clusters included TCRs from different participants, underscoring their public nature, and indeed for 5 of the 7 epitopes that contained clones from >1 participant, we detected no difference between the distributions of intra-participant v inter-participant TCR distances (0.16<p<1 by Kolmogorov–Smirnov test), indicating that features recognizing the common peptide:HLA epitope often predominate over any participant-to-participant differences in TCR repertoires. A striking exception to this was the DRB1^*^15:03_HVSFVMAYPEMLAA epitope: whereas the TCRs from participants 40059 and 30129 clustered indistinguishably (red v blue dots in the left box of **Figure 6a**: p(Inter v Intra | 40059v30129) = 0.69), the TCRs from 30168 (green dots) were an outlier (p(Inter v Intra | 30168v40059) = 0.001; p(Inter v Intra | 30168v30129) = 0.0005), driven by a dearth of 30168 epitope-specific TCRs in the epitope’s central homology cluster, and a correspondingly greater frequency of non-clustered TCRs. Intriguingly, this outlier participant (30168) is a homozygote for the restricting allele (DRB1^*^15:03), and in fact represents the only case among the 7 DRB1-restricted epitopes for which the participant was homozygous.

Together, these findings indicate that public TCR sequence features arise in the response to all or most *Mtb* epitopes: occasionally (e.g. for the PE13 epitope) these are prominent and can characterize the majority of responding TCRs, however for most epitopes, sequence homologies are relatively rare and likely only detectable when the analysis is focused on TCRs from epitope-specific cells (e.g. by filtering on probe binding).

## Discussion

In this study, we present a platform – the Clonally-Resolved Epitope-Specificity and Transcriptome Assay (CRESTA) – that enables the integrated analysis of CD4 T cell specificities and phenotypes at a dimensionality that greatly exceeds what was previously possible. By applying this assay to study the response across 100s of HLA class II-restricted epitopes in *Mtb*-infected participants, we generate a portrait of CD4 T cell immunity to *Mtb* at unprecedented breadth and resolution. This analysis reveals previously undescribed features including extensive intra-epitope polyclonality and phenotypic heterogeneity, both within and between epitopes. In the process, we also map 100s of new TCRα:β pairs to *Mtb* peptide:HLA epitopes, and describe how public epitope-specific sequence features vary across epitopes and individuals. More generally, since its assay targets are fully customizable, we expect CRESTA to be broadly applicable beyond *Mtb*, and to enable similar insights in other research/disease settings in which antigen-specific CD4 T cell responses are implicated.

Our finding that human *Mtb* epitope-specific T cells can occupy a range of differentiation states – including Th17, Treg, and a range of Th1 substates – within the same host, and sometimes even against the same epitope (**Figures 4, 5**) – illuminates layers of diversity in this response that were not previously understood. Especially notable is the related observation that different epitope specificities can elicit markedly different Th1 phenotypes within the same response (**Figure 5b**,**c**). Together, these findings indicate that existing approaches for studying *Mtb*-specific T cell responses – which are typically characterized by the use of antigen pools and the detection of a limited number of phenotypic markers (e.g. IFNγ, TNF)^28,36^ – capture only a subset of the response. The finding that different *Mtb* antigens can elicit T cells with different phenotypes/functions also offers a new class of mechanism to inform the interpretation of prior results that point to antigen-specific protection, including: (i) that different *Mtb* antigens are under purifying v diversifying evolutionary selection pressure^37^, (ii) that particular TCR clusters (i.e. epitope-specific responses) are associated with protection v non-protection from disease^31^. Particularly intriguing is our finding that the same PE13 epitope that was associated with protection in the latter study, can be uniquely associated with high expression of GNLY (**Figure 5c**), a pore-forming effector found in cytotoxic granules that can kill *Mtb* bacteria at high concentrations^38,39^. More generally, our findings invite future applications of CRESTA to larger cohorts of *Mtb*-infected participants, including comparisons of progressors v controllers, to explore more comprehensively which T cell epitope-specificities and phenotypes may be associated with protection from disease.

Our unbiased, epitope-resolved analysis of >200 TCRα:β sequences identified by CRESTA (**Figure 5**) yields several important insights. First is the finding that, when filtering stringently on TCRα:β pairs specific for individual *Mtb* epitopes, some degree of homology is detectable for all epitopes. This observation is consistent with the first-principles consideration that, although most epitopes are likely capable of recognition by many possible TCR binding modes, each binding sequence is likely to be surrounded by a cluster of similar sequences separated by amino acid substitutions that preserve binding. Second, however, is the countervailing observation that, even in our stringent setting (filtering on individual epitopes), these homologies encompass only a minority of antigen-specific TCRs. Moreover, when extrapolating to the more typical scenario where oligoclonal/polyclonal repertoires are analyzed, such homologies remained detectable in only rare cases – implying a limit to the sensitivity of approaches that rely on TCR clustering alone. Third, although the most common finding was indistinguishable TCR similarity patterns within v between individuals, our observation of a striking exception – in which one participant showed dramatically weaker TCR clustering for the PE13 epitope – represents the best evidence of which we are aware that there can be large individual-specific differences in the TCR sequences raised to a fixed peptide:HLA antigen. Conceivable interpretations include: (i) individual-to-individual differences in the T cell repertoire, shaped by other HLAs, self-peptides and/or antigen exposures, (ii) individual-to-individual differences in the milieu in which the epitope-specific T cells were primed, potentially altering the affinity threshold for T cell activation, (iii) a gene-dose effect of the restricting HLA, wherein homozygosity may increase the density of peptide:HLA complexes and thereby decrease the affinity threshold for T cell activation. Finally, CRESTA’s ability to generate large numbers of TCRα:β sequences mapped to individual peptide:HLA epitopes (as of June 2024, considering HLA class II:peptide-specific TCRα:β pairs in the IEDB, this study alone generated ∼8X more pairs for *Mtb*, and >10% of the total number of pairs identified across all research fields), positions it as a powerful tool for TCR discovery, with the potential future applications that include the development of TCR-based therapies, as well as the training of next-generation models for predicting antigen specificity from TCR sequences.

A cornerstone of CRESTA is clonal analysis, which we use to (i) make rare epitope-specific CD4 T cells detectable within limited sample volumes, and (ii) enable robust, multi-datapoint-based inferences about their antigen specificity and phenotype from single cell sequencing data. At the same time, the use of clonal expansion has the potential to limit the assay in several ways. First, it is likely that the efficiency of such expansion varies according to the state of the precursor T cells, meaning that highly proliferative subtypes may reach detectable levels before less proliferative subtypes do, biasing the representation of cells analyzed. Indeed, within cells expanded from *Mtb*-infected participants, we observe a significant correlation between clone size and helper T cell state (**Figure 4c**). Nonetheless, CRESTA successfully detects cells across the Th state spectrum, including numerous clones of the less proliferative Treg and Th2 subtypes (**Figure 4**), and its sensitivity to less expanded clones is likely to increase as the throughput of single cell sequencing technologies continues to grow. Importantly, the impact of such biases can be controlled by comparing clones of interest against bystander clones within the same expanded samples, as we exemplify in **Figure 4d**. It is also noteworthy that the sensitivity of common (non-expansion-based) T cell specificity assays is often also biased to particular phenotypes (eg ELISpot, that depends on cytokine secretion). A second potential limitation of clonal expansion is the possibility of introducing artefactual gene expression patterns. While we have not directly quantified these effects, our observation of gene expression patterns that partition strongly according to clonal identity within a mixed culture (**Figures 2d,e**) – and recapitulate the known Th subsets upon unsupervised multi-gene analyses (**Figure 4a,b**) – indicates that initial differentiation states persist during expansion to at least a substantial degree. In the future, any remaining impact may be mitigated by reducing the degree of expansion (our observed clone size distributions indicate that even an order of magnitude less expansion is unlikely to substantially impact assay sensitivity), and/or by using other sample types (e.g. bronchoalveolar lavage) that are more enriched for the relevant antigen-specificities.

While it represents a major advance beyond what has been enabled in prior studies of CD4 T cells, the application of CRESTA at a multiplexity of up to ∼170 epitopes in this work does not yet reach the scale that is needed for broad proteome-scale epitope discovery and characterization. However, since: (i) we expect DNA barcoding and sequencing to be intrinsically highly-scalable (>100,000s-plex), (ii) we saw no evidence for loss of binding signal as plexity increased up to ∼170 (**Figure 3e**), and (iii) analogous approaches for CD8 T cells been successfully implemented up to ∼1000-plex (even without the clonal analysis enhancement we describe here)^11^, we expect CRESTA to scale well beyond the plexity that we have demonstrated here. At that point, new bottlenecks to be overcome will be the efficient identification of candidate peptide:HLA pairs (e.g. using *in silico* prediction and/or high-throughput binding screens), and the preparation of the corresponding probes in large numbers (e.g. using microfluidic automation). Realizing these advances could enable powerful new studies of CD4 T cell specificities and functions, and their interactions, at a truly genome-wide scale in diverse disease states.

## Methods

### Study cohort, PBMC collection and processing

Household contacts (HHCs) of newly diagnosed active pulmonary TB cases were referred to the Kenya Medical Research Institute (KEMRI) Clinical Research Center in Kisumu, Kenya, and their demographic and medical history data were collected. HHCs were persons who shared the same home residence as the index case for ≥5 nights during the 30 days prior to the date of TB diagnosis of the index case, and were enrolled no more than 3 months (mean: 18 days; range: 1–77 days) after the index case began TB treatment. All participants provided written informed consent to join the study and were recruited from two community-based health clinics located in Kisumu City and Kombewa, Kisumu County. All enrolled individuals met the following inclusion criteria: ≥ 13 years of age at the time of enrollment, positive QuantiFERON TB Gold in Tube (QFT) result, seronegative for HIV antibodies, no previous history of diagnosis or treatment for active TB disease or LTBI, normal chest X-ray, and not pregnant. All participants were presumed to be BCG vaccinated due to the Kenyan policy of BCG vaccination at birth and high BCG coverage rates throughout Kenya. All participants gave written informed consent for the study, which was approved by the KEMRI/CDC Scientific and Ethics Review Unit and the Institutional Review Board at Emory University, USA.

Blood samples were collected from participants in sodium heparin or lithium heparin Vacutainer CPT Mononuclear Cell Preparation Tubes (BD Biosciences or Greiner Bio-One). PBMC were isolated by density centrifugation, rested in complete media (RPMI 1640 containing L-glutamine supplemented with 10% heat-inactivated fetal bovine serum (FBS), 1% PenStrep, 1% Hepes) before counting. PBMC isolation was initiated <2 hours after the blood was drawn. Isolated PBMC were cryopreserved in 90% heat-inactivated fetal calf serum/10% DMSO, and kept in LN2 (and shipped on dry ice) until they were thawed for study at the TGen laboratory.

### Cell culture and T cell expansion

PBMCs were thawed in a 37°C water bath and then washed and plated at 1.25 × 10^6^ cell/mL in 24-well flat-bottomed plates in complete RPMI (RPMI-1640 with 10% AB human serum, 0.8 mM sodium pyruvate, 0.8x non-essential amino acids, 80 U/mL penicillin-streptomycin, 0.4x HEPES, 200 mM L-glutamine, 0.07x 2-Mercaptoethanol; hereafter “cRPMI”) at 37°C with 5% CO_2_. On day 2, pools of peptides dissolved in DMSO (up to 579 peptides – which included all peptides from multimers used for staining – each at a final concentration 0.47ug/mL; with a maximum final DMSO concentration of 1.3%) were added. On day 3, taking care to not disturb the cells, 50% of the media was exchanged. Cells were split as needed on days 4-5, after which 50% of the media was again exchanged. Cultures were maintained for a total of 8-10 days. Media included recombinant human interleukin-2 (IL-2) at 230IU/mL – 1,025IU/mL (Biolegend), with the lower concentrations used on days 1-2, followed by higher concentrations throughout the remainder of the expansion. At the end of the culture, cells were harvested by collecting the supernatant, treating wells with 2 mM EDTA in PBS for 2-4 minutes and adding the detached cell suspension to the collected culture. After combining all wells from the same donor, cells were spun down at 300 x g for 5 min at room temperature, resuspended in storage media (9:1 cRPMI and DMSO) and stored in liquid nitrogen until further use, or resuspended in cRPMI and used directly in the CRESTA assay.

### Production of HLA:peptide complexes

Peptides were custom ordered from Millipore-Sigma (PEPscreen, unpurified) and reconstituted to 20 mg/mL in DMSO. CLIP peptide-tethered, biotinylated HLA monomers were obtained from the NIH Tetramer Core Facility. To generate peptide-bound HLA monomers, we used a protocol described previously^32^, consisting of (i) CLIP peptide cleavage, followed by (ii) peptide exchange. CLIP peptide cleavage was performed by incubating the HLA monomer (at a final concentration of 0.4 mg/mL with 3C protease (HRV-3C protease, Sigma Aldrich) in 3C cleavage buffer (0.05 M Tris pH, 7.5, 150 mM NaCl) or thrombin protease in 10X Thrombin cleavage buffer (Thrombin restriction grade, Millipore) overnight at room temperature. To perform peptide exchange, the cleaved monomer was incubated at a final concentration of 0.2 mg/mL with individual peptides (at a final concentration of 76-666 ug/mL) in 50 mM citrate buffer pH 5.2 containing 100 mM NaCl, 2 mM EDTA, 0.2x protease inhibitor (Promega), at reaction scales of 30–1000 μL depending on desired yield, at 30°C for 4 days. Following incubation, HLA:peptide complexes were cleared of excess peptide and concentrated using dPBS-rinsed Amicon Ultra-0.5 Centrifugal 10 kDa filters (spun twice at 14,000 x g for 15 min at 4°C). Finally, the cleared HLA:peptide monomer products were quantified using a Nanodrop 1000 spectrophotometer reading at 280 nm, and stored in the presence of 0.75x protease inhibitor cocktail (50X protease inhibitor, Promega) at -80°C for up to 12 months.

### Generation of DNA-barcoded HLA:peptide probes

Barcoding DNA sequences compatible with the 10X Chromium Single Cell 5’ chemistry were designed according to the 69 mer construct recommended by 10X Genomics (Surface Protein Labeling Protocol CG000186), utilizing a 3’ Capture Sequence and containing 15 mer barcodes from the 10X barcodes whitelist (Demonstrated Protocol, CG000193). These sequences were purchased from IDT as 5’ biotinylated DNA oligos with standard desalting. For each desired multimer probe, 1 μL of streptavidin-bearing dextran backbone (Klickmer APC or PE, Immunodex, 0.16 μM) was barcoded by incubating it with 1 μL of barcoding oligonucleotide (at 0.32 μM in 1x TE) at 4°C for 30 min, in Lo-bind (Eppendorf) 96 well plate (stoichiometry = 1 dextran : 2 oligos). After incubation, 2 μL of the desired HLA:peptide monomer (at 3.2 μM, prepared as above) was added to the barcoded dextramer and incubated for 30 min at RT (stoichiometry = 1 dextran : 2 oligos : 20 HLA:peptide monomers). Binding reactions were quenched by adding 1 μL free D-biotin (5μM) in excess at 4°C for 30 min. These individual probe constructs (“multimers”) were stored for up to 1 week at 4°C prior to cell staining. On the day of cell staining, the total volume (5 μL) of each desired multimer (up to 173 – see **Supplementary Table 2** for the probes used for each donor) was pooled and concentrated using a Vivaspin2 column (#VS0241; Sartorius). The column was pre-washed with 1x PBS, followed by 2 mL of barcode buffer (dPBS with 0.5% BSA and 5 μg/mL Herring DNA (Promega)) and then stored at 4°C prior to loading with the probe pool. Once the probe pool was added, the column was spun at 3,300 rpm for 10–20 minutes (until the liquid level reached ∼50-100 μl), and then inverted and spun at 2,000 x g for 5 minutes to recover the pool, which was kept on ice until cells were ready for staining (up to 2 hours).

### PMA/ionomycin stimulation (“gene expression aliquot”)

Expanded cells were thawed, washed with cRPMI, and then resuspended in cRPMI with 200 IU/mL IL-2 at 1.25 × 10^6^ cell/mL in 24-well flat-bottomed plates and incubated at 37°C with 5% CO_2_ overnight. Cells were then stimulated with PMA and ionomycin at final concentrations of 0.04 μM and 0.67 μM, respectively (Biolegend, Catalog #423301), and then incubated for 1-1.5hrs at 37°C. Stimulated cells were collected into 1 5mL conical tubes, including detachment from plates with 2 mM EDTA in PBS for 2-4 minutes, then centrifuged at 200 x g for 5 minutes. Cells were washed a total of three times, in the following buffers, with spinning at 300 x g for 5 min at RT between each: (i) 5 mL of cRPMI; (ii) 2.5 mL cRPMI + 2.5 mL EasySep Buffer (# 20144; StemCell); and (iii) 5 mL of Loading Buffer (1x PBS + 0.04% BSA). Following the final wash, cells were resuspended in 100μL of Loading Buffer and a 10 μL aliquot was taken for cell counting prior to single cell partitioning.

### DNA-barcoded peptide:HLA class II multimer staining (“antigen-specificity aliquot”)

Expanded cells were thawed and rested overnight as described above (*PMA/ionomycin stimulation* section). The next day, cells were collected into 15 mL conical tubes, including detachment from plates with 2 mM EDTA in PBS for 2-4 minutes, then centrifuged at 200 x g for 5 minutes. Supernatant was discarded without disturbing the cell pellet, and cells were washed 2 times in cRPMI (300 x g for 5 min) and the pellet resuspended (via flicking) in 100 μL cRPMI. Resuspended cells treated with 0.003 M of the protein kinase inhibitor Dasatinib (Axon Medchem, VA) with gentle swirling at 37°C, for 10 min with a loose cap, and then 20 min with a tight cap. The multimer probe pool (50-100 uL total volume, prepared as above) was then added, resulting in a final concentration of ∼0.8–1.1 nM for each multimer, and cells incubated for 30 min at RT with a tight cap. After incubation, cells were spun in a 4°C (300 x g for 5 min) for 3 washes; first in 5 mL of cRPMI, then 2.5 mL cRPMI + 2.5 mL EaspSep Buffer (# 20144; StemCell) and finally in 5mL Loading Buffer (1x PBS + 0.04% BSA). Following the last wash, cells were resuspended in 100 μL Loading Buffer and a 10 μL aliquot was taken for cell counting prior to single cell partitioning.

### Generation and sequencing of single cell libraries

To generate single cell libraries, 10,000–40,000 cells per participant aliquot were loaded into the Chromium Controller (10X Genomics), and processed according to the manufacturer’s instructions (Chromium Next GEM Single Cell 5’ Reagent Kits v2 protocol) to prepare libraries corresponding to the feature/probe barcodes (antigen-specificity aliquot), 5′ gene expression (gene expression aliquot), and VDJ-T repertoire (both aliquots). Fragment size and quantity was assessed on the 4200 TapeStation (#G2991A; Agilent) using high sensitivity DNA tapes (#5067– 4626; Agilent). Before sequencing, qPCR was performed after serial dilution of the libraries (#KK4824 – 07960140001; Kapa Biosystems). Libraries were sequenced on Illumina NextSeq1000 or NovaSeq instruments using the read configurations and PhiX loading recommended by 10X Genomics.

### Analysis of single cell sequencing data

#### Data processing and merging

Following Illumina BCL file conversion, fastq files were processed through the CellRanger *Multi* Pipeline (10X Genomics) to perform demultiplexing, alignment, filtering, barcode and UMI counting, and VDJ assembly. We then developed custom R code for all downstream analytical steps. We first merged the filtered VDJ clonotype sequences (from the *filtered_contig_annotations*.*csv* file) with the raw counts matrix of transcripts and/or probe barcodes (*raw_feature_bc_matrix*), by matching on cell barcodes and retaining only those cells associated with ≥1 filtered VDJ sequence.

#### Clonotype collapsing and shortlisting

Since some CellRanger VDJ-T clonotype calls contain the same TCR chain sequence both alone and paired with other sequence(s) (likely attributable to a combination of: (i) droplets containing >1 cell (multiplets), and (ii) the fact that TCR chain sequences that are present do not always successfully amplify/assemble in every cell), we developed a method to improve assignment of TCR chains to clonotypes that involved clustering chains according to their correlated occurrence across cells (*collapseclonotypes*.*R*). This allowed us to more accurately identify chains belonging to the same clone, by merging cells that were assigned to different clonotypes but shared ≥1 chain, and by excluding cells that contained chains from ≥1 cluster (as likely multiplets). For each aliquot (antigen-specificitiy or gene expression) we then focused only on collapsed clones containing ≥3 cells (*listclonotypes*.*R*).

#### Normalization of multimer probe signal

We developed a method to normalize multimer-to-multimer and cell-to-cell differences in the multimer assay yield *(normmaster*.*R)*. First, we normalized the read counts for each multimer to a fixed depth such that the sums for all multimers were the same (equal to the number of cells in the data matrix). For each multimer, we then normalized in the other dimension (across cells) by dividing the resulting values for each cell by the median value of all other cells. To reduce the impact of high-staining outliers (e.g. caused by multimer aggregation) on visualization, for each multimer we applied the value of the cell at the top 0.1 percentile to all cells with greater staining. Finally, for each multimer, we divided values by the multimer max value, to scale values onto the [0,1] interval prior to visualization (*stackedplot*.*R*) and statistical analysis.

#### Identification of staining multimers / clonotypes and Clonal Differentiation Genes

To identify binding multimers and Clonal Differentiation Genes (CDGs), we applied Kruskal-Wallis tests to quantify the influence of clonal identity on (i) the normalized multimer probe signal, or (ii) raw reads, for each multimer or gene, respectively (*testclonalgenes*.*R*). To identify the individual binding clones for each of binding epitope, we then applied one-tailed Wilcoxon tests post-hoc to each binding multimer, in which we compared the normalized multimer probe signals in each individual clone against the signal of the central 20% of cells across all clones (*testsigclones*.*R*).

#### UMAP analysis, transcript correlation analysis and identification of T cell subsets

Beginning with either all clones (**Figure 4**) or all epitope-mapped clones (**Figure 5c**), we identified significant CDGs as above (with max fold-difference > 10 and Bonferroni-adjusted p-values < 0.05), and then generated log_10_(median+1) values (“clone medians”) for each gene in each clone. To the resulting gene-by-clone table, we applied UMAP to generate a 2-dimensional rendering using the R *umap* package. For the lineage defining genes IFNγ, IL4 and IL17A, we generated Pearson correlation coefficients (r) across clone median values, against every other significant CDG. Genes with r > 0.5 are depicted as a force-directed graph rendered using the R *igraph* package. Using clone median values, we identified Th2, Th17 and Treg clones as IL4+GATA3+, IL17A+RORC+, and FOXP3+CTLA4+, respectively. Among the remainder, we identified Th1 clones as IFNG+CCL4+, and finally, among the remainder, naives clones as IL2RA–.

#### Analysis of TCRα:β homologies using TCRdist

To analyze TCR homologies, we developed our own computationally efficient implementation of the TCRdist metric^35^ to quantify the degree of sequence similarity between pairs of TCRα:β heterodimers. We focused on epitope-specific clones with exactly 1 TCRα and 1 TCRβ chain, and, within each epitope, performed TCRdist measurements between all pairs of binding TCRs. We then mapped each TCRdist value to a p-value using a large distribution of TCRdists measured on a large, unenriched repertoire, as follows. Beginning with 1e4 randomly-sampled TCRαs and TCRβs, we calculated all ∼5e7 pairwise TCRdists for each chain type. We then calculated the frequencies of all possible α:β chain TCRdist values by considering all combinations of α and β TCRdists, allowing us to efficiently estimate frequencies down to ∼1e-12. Epitope-specific sets of p-values were adjusted using the Benjamini–Hochberg procedure. We visualized the CDR3 sequences of significantly clustered TCRs using the *ggseqlogo* R package, after multiple sequence alignment. Positions in which there was an alignment gap for the majority of CDR3s were excluded from display.

